# The PH/MyTH4/FERM molecule MAX-1 inhibits UNC-5 activity in regulation of VD growth cone protrusion in *Caenorhabditis elegans*

**DOI:** 10.1101/2021.08.25.457713

**Authors:** Snehal S. Mahadik, Erik A. Lundquist

## Abstract

UNC-6/Netrin is a secreted conserved guidance cue that regulates dorsal-ventral axon guidance of *C. elegans* and in the vertebral spinal cord. In the polarity/protrusion model of VD growth cone guidance away from ventrally-expressed UNC-6 (repulsion), UNC-6 first polarizes the growth cone via the UNC-5 receptor such that filopodial protrusions are biased dorsally. UNC-6 then regulates a balance of protrusion in the growth cone based upon this polarity. UNC-5 inhibits protrusion ventrally, and the UNC-6 receptor UNC-40/DCC stimulates protrusion dorsally, resulting in net dorsal growth cone outgrowth. UNC-5 inhibits protrusion through the flavin monooxygenases FMO-1, 4, and 5 and possible actin destabilization, and inhibits pro-protrusive microtubule entry into the growth cone utilizing UNC-33/CRMP. The PH/MyTH4/FERM myosin-like protein was previously shown to act with UNC-5 in VD axon guidance utilizing axon guidance endpoint analysis. Here, we analyzed the effects of MAX-1 on VD growth cone morphology during outgrowth. We found that *max-1* mutant growth cones were smaller and less protrusive than wild-type, the opposite of the *unc-5* mutant phenotype. Furthermore, genetic interactions suggest that MAX-1 might normally inhibit UNC-5 activity, such that in a *max-1* mutant growth cone, UNC-5 is overactive. Our results, combined with previous studies suggesting that MAX-1 might regulate UNC-5 levels in the cell or plasma membrane localization, suggest that MAX-1 attenuates UNC-5 signaling by regulating UNC-5 stability or trafficking. In summary, in the context of growth cone protrusion, MAX-1 inhibits UNC-5, demonstrating the mechanistic insight that can be gained by analyzing growth cones during outgrowth in addition to axon guidance endpoint analysis.

## Introduction

Many guidance cues and receptors that control directed cell and growth cone migrations have been identified, but the cellular mechanisms by which they influence migration *in vivo* remain to be fully elucidated. UNC-6/Netrin is a conserved secreted guidance cue that regulates dorsal-ventral axon guidance in metazoans, including *C. elegans* and the vertebral spinal cord (Hedgecock *et al*. 1990; Ishii *et al*. 1992; Leung-Hagesteijn *et al*. 1992; Chan *et al*. 1996; Leonardo *et al*. 1997; Hong *et al*. 1999; Montell 1999; Shekarabi and Kennedy 2002; Moore *et al*. 2007). In *C. elegans*, UNC-6/Netrin is required for both dorsal and ventral axon outgrowth. UNC-6/Netrin is expressed in ventral cord motor neurons (Wadsworth *et al*. 1996). Dorsally-directed axons are repelled from UNC-6/Netrin, and ventrally directed axons attracted to it. Classically, Netrin was thought to establish a ventral-to-dorsal gradient, which was followed by growth cones growing up the gradient (attraction) or down the gradient (repulsion) (Serafini *et al*. 1996; Fazeli *et al*. 1997; Kennedy *et al*. 2006). Recent studies in vertebrate spinal cord and our studies in *C. elegans* suggest a distinct mechanism (Dominici *et al*. 2017; Varadarajan *et al*. 2017; Yamauchi *et al*. 2017; Morales 2018). In the spinal cord, floorplate Netrin1 was found to be dispensable for ventral commissural axon guidance. Rather, ventricular zone Netrin1 expression was required, suggesting a close-range, haptotactic mechanism. In *C. elegans*, we found that UNC-6/Netrin controls growth cone polarity and growth cone protrusion (Norris and Lundquist 2011; Norris *et al*. 2014; Gujar *et al*. 2018; Gujar *et al*. 2019). In the VD motor neuron growth cones that grow dorsally away from the ventral nerve cord, UNC-6/Netrin and its receptor UNC-5 polarize the growth cone in the dorsal-ventral axis, such that growth cone filopodial protrusions are dorsally-biased. Based on this polarity, UNC-5 then inhibits growth cone protrusion ventrally in response to UNC-6/Netrin. The UNC-6/Netrin receptor UNC-40/DCC stimulates protrusion dorsally. This polarity/protrusion model involves asymmetry of receptor function across the growth cone in response to UNC-6/Netrin, rather than a gradient. Similar regulation of polarization and protrusion was observed in growth cones attracted to UNC-6/Netrin (the Statistically-Oriented Asymmetric Localization (SOAL) model) (Limerick *et al*. 2017).

UNC-6 mediates a balance of pro- and anti-protrusive forces in the VD growth cone required for proper outgrowth (Norris and Lundquist 2011; Gujar *et al*. 2018). In *unc-5* mutants, VD growth cones are overly-protrusive, with larger growth cone area and longer growth cone filopodia (Norris *et al*. 2014), indicating that UNC-5 inhibits protrusion. Indeed, constitutively-active MYR::UNC-5 results in small growth cones with reduced filopodial protrusion (Norris and Lundquist 2011). UNC-40 is required for the excess protrusion in *unc-5* mutants (Norris and Lundquist 2011), suggesting that pro-protrusive UNC-40 activity predominates in an *unc-5* mutant. Our previous work shows that UNC-5 inhibits protrusion through at least two distinct mechanisms. The flavin monooxygenases FMO-1, FMO-4, and FMO-5 act downstream of UNC-5 to inhibit protrusion (Gujar *et al*. 2017), possibly by destabilizing the actin cytoskeleton similar to MICAL (Terman *et al*. 2002). UNC-33/CRMP acts downstream of UNC-5 to inhibit protrusion and might act by restricting microtubule entry into the growth cone (Norris *et al*. 2014), which are pro-protrusive (Gujar *et al*. 2018).

Previous studies showed that the PH/MyTH4/FERM myosin-like molecule MAX-1 (motor axon guidance-1) acts in the UNC-5 pathway in parallel to UNC-40 in VD/DD commissural axon guidance (Huang *et al*. 2002). Furthermore, MAX-1 interacts with the cytoplasmic region of UNC-5 until it is sUMOylated, at which point it is displaced by the ABP-3/AP-3 complex β molecule, which targets UNC-5 to the lysosome for degradation (Chen *et al*. 2018). Thus, MAX-1 might control UNC-5 levels in the cell. In support of this idea, a study in zebrafish suggested that *max-1* plays a role in regulating membrane localization of Ephrin3b protein, which provides guidance cue for the migration of endothelial cells during embryogenesis (Zhong *et al*. 2006). In this model, MAX-1 is constitutively bound to UNC-5 until it is SUMOylated, which allows ABP-3 to bind to UNC-5 and target it for lysosomal degradation (Chen *et al*. 2018). These previous *C. elegans* studies relied on endpoint axon guidance analysis and did not assess the effect of MAX-1 on growth cones during outgrowth.

In this study, we analyzed VD growth cones during outgrowth in *max-1* mutants and in double mutants with *unc-5, unc-6*, and *unc-40* to understand the role of MAX-1 with regard to UNC-5 activity in the growth cone. We found that VD growth cones in *max-1* mutants were small and less-protrusive than wild-type, similar to constitutively-active *myr::unc-5. max-1* mutants did not show strong growth cone polarity defects, suggesting it might specifically affect protrusion. *unc-5* loss-of-function display the opposite phenotype, with larger and more protrusive growth cones, suggesting that UNC-5 and MAX-1 have opposing roles in growth cone protrusion. Double mutants of *max-1* with *unc-5* hypomorphic mutants, which retain *unc-5* function, displayed growth cones that resembled the small, less-protrusive growth cones of *max-1* alone. This suppression of *unc-5* hypomorphic mutants could be due to the fact that remaining UNC-5 activity in these mutants is no longer attenuated by MAX-1, and thus inhibits growth cone protrusion. This is consistent with a role of MAX-1 in controlling UNC-5 levels in the cell. Growth cone protrusion in *unc-40; max-1* and *unc-6; max-1* double mutants did not suggest strong genetic interactions and are consistent with MAX-1 acting in the UNC-6 pathway. However, endpoint axon guidance analysis confirmed that MAX-1 acts in parallel to UNC-40.

In sum, our results suggest that MAX-1 normally inhibits UNC-5 activity in the growth cone, possibly by regulating UNC-5 levels in the cell or at the plasma membrane. This is in contrast to these earlier studies which suggested that MAX-1 might stimulate or mediate UNC-5 activity. Clearly, there is more to learn about the complex activity of MAX-1 and UNC-5 in the growth cone. In any case, analysis of growth cone morphology during outgrowth can add mechanistic detail to the complex roles of axon guidance molecules and a deeper understanding of how growth cone outgrowth results in proper axon guidance and circuit formation.

## Material and Methods

### Genetic Methods

Experiments were performed at 20^0^C using standard *C. elegans* techniques (Brenner 1974). Mutations were used LGI: *unc-40(n324);* LGII: *juls76 [Punc-25::gfp];* LGIV: *unc-5(e791, ev480* and, *e152*); LGV: *max-1(ju39, ju142 and lq148)*; LGX: *unc-6(ev400 and e78)*. The presence of mutations was confirmed by phenotype and sequencing. Chromosomal locations not determined. *myr::unc-5* [*Punc-25::myr::unc-5::gfp*]. *max-1(lq148)* was identified by whole genome sequencing after EMS mutagenesis. *lq148* is a C to T transition that creates a glutamine to stop codon at codon 857 (position 13,099,591 on LGV). *lq148* failed to complement *max-1(ju39)* for the Unc phenotype and VD/DD axon guidance defects (data not shown).

### Imaging of Axon guidance defects

VD/DD neuron were visualized with *Punc-25::gfp* transgene, *juIs76* (Jin *et al*. 1999), which is expressed in 19 GABAergic commissural motor neurons including DD1-6 and VD1-13. The commissure on the left side (VD1) was not scored. The remaining 18 commissures extend processes on the right side of the animal. Due to the fasciculation of some commissural processes, an average 16 VD/DD commissural processes were apparent in wild-type. Total axon guidance defects were calculated by counting the number of commissural processes that failed to reach the dorsal nerve cord, wandered laterally at an angle of 45^0^ before reaching the dorsal nerve cord, and ectopic branching (Gujar *et al*. 2017). Significance of difference between genotypes was determined using Fisher’s exact test.

### Growth cone imaging and quantification

VD growth cones were imaged as previously described (Norris and Lundquist 2011; Norris *et al*. 2014; Gujar *et al*. 2017; Gujar *et al*. 2018; Gujar *et al*. 2019). Briefly, wild-type animals were harvested 16-hour post-hatching at 20^0^C and placed on 2% agarose pads with 5mM sodium azide in M9 buffer. For mutant animals, due to the slower development the time point ranges between 17-21hour post hatching. Growth cone were imaged with Qimagine Rolera mGi camera on a Leica DM5500 microscope. Projections less than 0.5μm in width were scored as filopodia. Growth cone area and filopodial length were quantified using ImageJ software. Quantification was done as described previously (Norris and Lundquist 2011; Norris *et al*. 2014; Gujar *et al*. 2017; Gujar *et al*. 2018; Gujar *et al*. 2019). Significance of difference between two genotypes was determined by two sided *t*-tests with unequal variance.

Polarity of growth cone filopodial protrusion was determined as previously described (Norris and Lundquist 2011; Norris *et al*. 2014; Gujar *et al*. 2017; Gujar *et al*. 2018; Gujar *et al*. 2019). Briefly, growth cones were imaged as above, and divided into dorsal and ventral halves relative to the ventral nerve cord. Filopodial protrusions from the dorsal and ventral halves were counted and expressed as a percentage of dorsal protrusions (# dorsal protrusions/# all protrusions). Significance of difference between genotypes was determined using Fisher’s exact test.

Strains and plasmids are available upon request. The authors affirm that all data necessary for confirming the conclusions of the article are present within the article, figures, and tables.

## Results

### MAX-1 is required for robust VD growth cone protrusion

MAX-1 is a member of the PH/MyTH4/FERM myosin-like family and contains an N-terminal coiled-coil region, followed by two pleckstrin homology domains (PH), a myosin tail homology region, and a FERM domain (Figure 1A). *max-1* alleles were first identified in a screen for abnormal synapse formation and axon guidance defects in the GABAergic VD/DD motor neurons (Huang *et al*. 2002). *max-1(ju39)* is a 28-nucleotide deletion within the second exon which cause a frameshift after amino acid 58, *max-1(ju142)* is an 81-nucleotide deletion at the exon/intron boundary of the fourth exon (Huang *et al*. 2002), and *max-1(lq148)* is a C to T transition in the penultimate *max-1* exon resulting in a premature stop codon at glutamine 857 (Figure 1A and B). All three alleles are predicted to be strong loss-of-function or null alleles.

**Figure 1.**
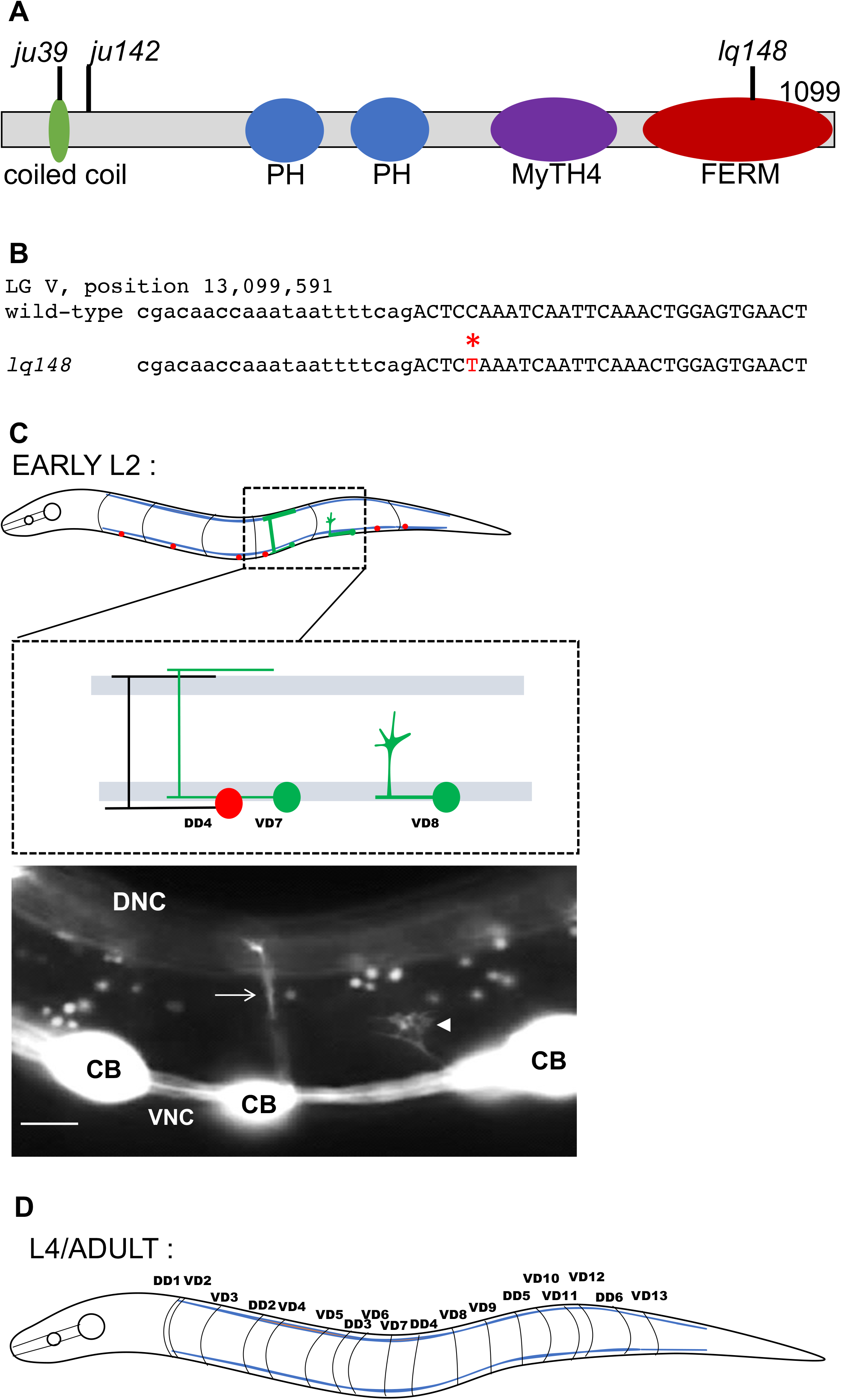
MAX-1 and VD/DD axon guidance. **(A)** The MAX-1 molecule with functional domains indicated. PH = pleckstrin homology domain. The locations of *max-1* mutants are indicted. **(B)** Sequence comparison of *lq148* allele to the wild-type. *lq148* shows change from C to T shown in red, at position 13,099591 on LG V, resulting in a premature stop codon at glutamine 857. **(C)** Diagram of an early L2 larva (Dorsal is up, and anterior is left). The blue lines represent the dorsal and ventral body wall muscle quadrants. The DD cell bodies (red) and axons (black) are shown. VD7 and VD8 are also shown (green). While migrating toward the dorsal nerve cord, VD8 exhibits a growth cone with dorsally-polarized protrusive morphology and dynamic filopodial protrusions. VD7 indicates the final structure of VD neurons in wild-type. The fluorescent micrograph shows a DD commissure (arrow) and a VD growth cone (arrowhead). The growth cone displays dorsally-directed filopodial protrusions. CB, cell body; DNC, dorsal nerve cord; VNC, ventral nerve cord. Scale bar represents 5μm. **(D)** Diagram of an L4 hermaphrodite after VD/DD axon outgrowth is complete. VD1 is not shown. The 18 right-side commissures shown. Some commissural axons extend as a single fascicle (e.g. DD1 and VD2) resulting in an average of 16 commissures per wild-type animal.

MAX-1 was previously shown to act with the UNC-6/Netrin receptor UNC-5 in dorsal VD/DD motor neuron axon guidance, and in parallel to the UNC-40/DCC Unc-6/Netrin receptor (Huang *et al*. 2002). These studies involved endpoint axon guidance analysis, but did not assay the growth cone during outgrowth. At the growth cone level, UNC-5 controls polarity of GABAergic growth cone protrusion as well as extent of protrusion (the polarity/protrusion model) (Norris and Lundquist 2011; Norris *et al*. 2014; Gujar *et al*. 2018). Given the genetic interaction of *max-1* with *unc-5*, we endeavored to understand the role of MAX-1 in GABAergic growth cone polarity and protrusion.

We first confirmed the GABAergic dorsal axon guidance defects in *max-1* mutants. The 19 VD/DD motor neurons cell bodies resides in the ventral nerve cord (Figure 1C). Axons of the VD and DD neurons first extend anteriorly in the ventral nerve cord, and then turn dorsally to extend commissural to the dorsal nerve cord (Figure 1C and Figure 2A). In wild-type animals, on average 16 commissures were observed, due to the fasciculation of some processes as single commissures (Figure 1D). As previously reported (Huang *et al*. 2002), *max-1* mutants display VD/DD axon guidance defects (21-25%), including axon wandering, failure to reach the dorsal nerve cord, and ectopic axon branching (Figure 2B-D and Figure 3).

**Figure 2.**
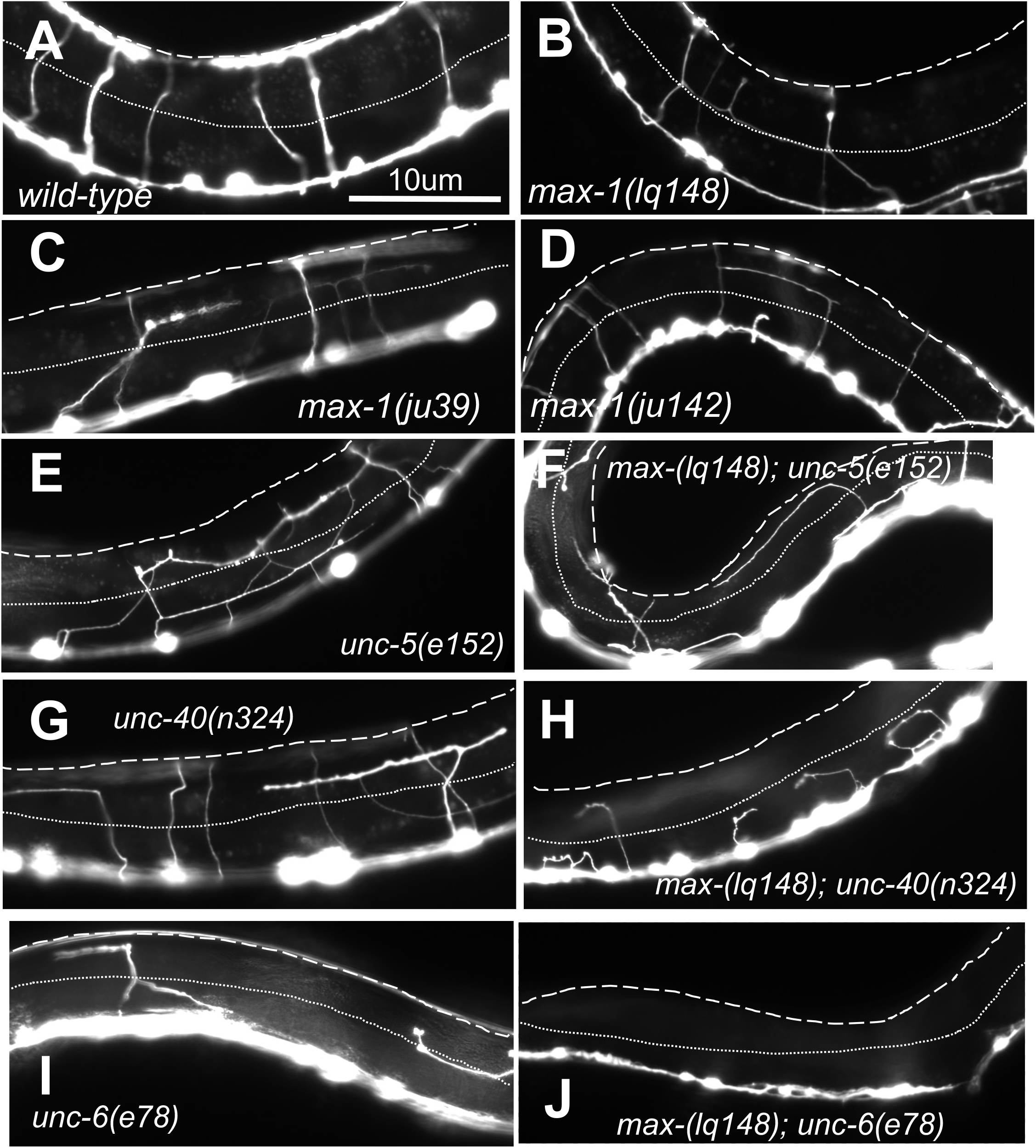
VD/DD endpoint axon guidance defects. Fluorescent micrographs of L4 animals of different genotypes are shown. Dorsal is up, and anterior is to the left. The scale bar in A represents 10μm for all micrographs. The dashed line indicates the dorsal nerve cord, and the dotted line the lateral midline.

**Figure 3.**
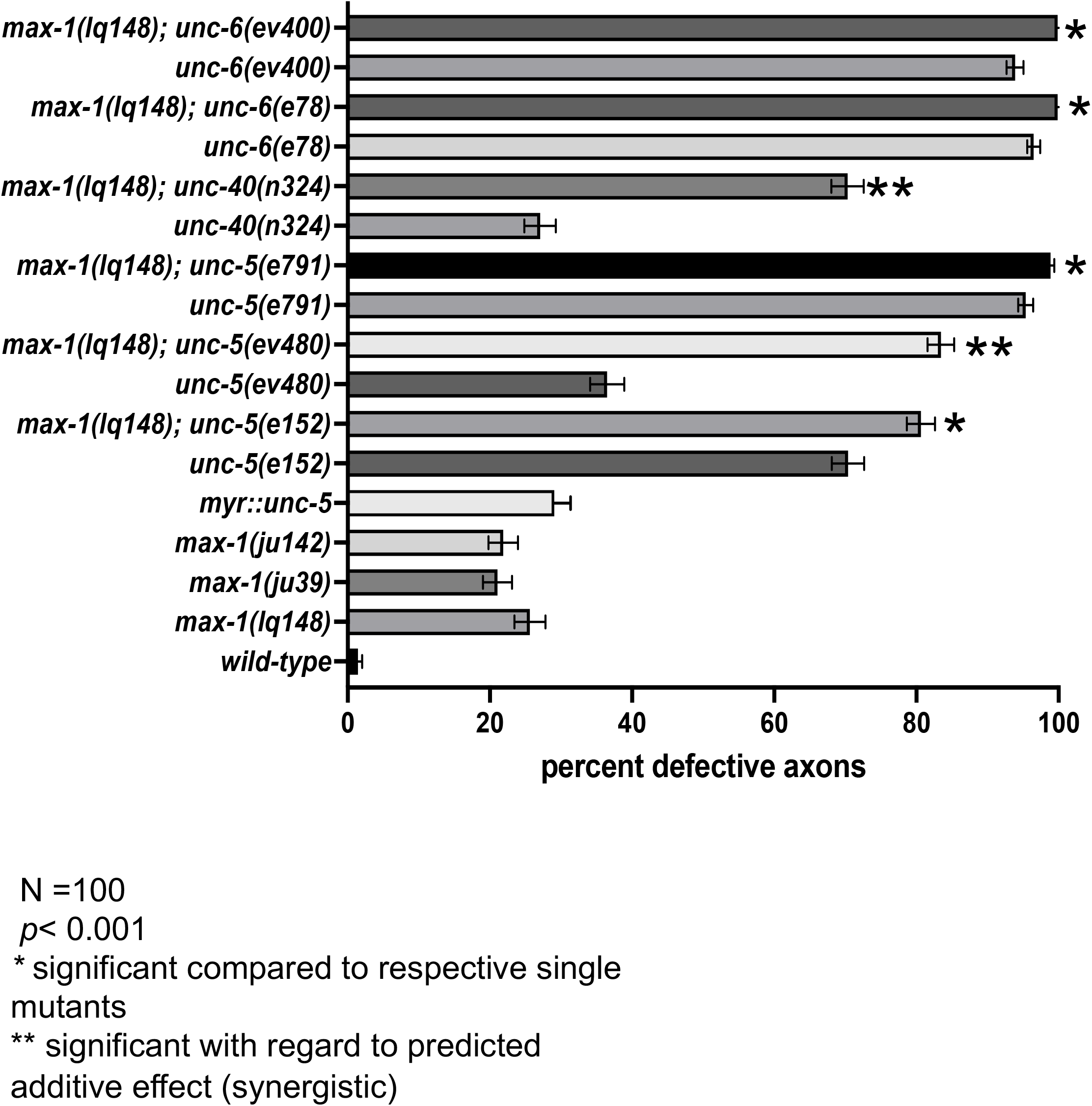
Quantification of VD/DD endpoint axon guidance defects. (A) Quantification of total axon guidance defects in wild-type and mutants (failure to reach the dorsal nerve cord, axon wandering, and branching). N = 100 animals/1600 commissural processes. Error bars represents 2X standard error of proportion; single asterisks (*) represents significance compared to respective single mutant; double asterisks (**) indicates a significant with regards to predicted additive effect (synergistic). The additive effect was calculated as p = p1 + p2 - (p1*p2). Fishers exact test was used to determine the *p* value (*p*< 0.001).

We next analyzed VD growth cone morphology in *max-1* mutants. In the early L2 larval stage, VD axons extend anteriorly in the ventral nerve cord, and then turn dorsally to extend a commissure to the dorsal nerve cord (Figure 1C). During this commissural extension, between the lateral midline and the ventral and dorsal nerve cords, wild-type VD growth cones display a robust growth cone body and filopodial protrusions (Figure 1C and Figure 4A, B, and D), with an average growth cone area of 4.6μm^2^, and an average filopodial length of 0.95μm (Figure 4A and B). Furthermore, filopodial protrusions were biased to the dorsal half of the growth cone (Figure 4C and D). In *max-1* mutants, VD growth cones were significantly smaller than wild-type, and filopodial protrusions were significantly shorter (2-3μm^2^ and 0.7-0.75μm, respectively; Figure 4A, B, and F-H). Filopodial protrusions were still biased to the dorsal, but less so than wild-type in all three alleles, and significantly less so in *ju142* (Figure 4C). In sum, these data indicate that MAX-1 is required for robust growth cone protrusion, with a lesser effect on growth cone polarity. As previously reported (Norris *et al*. 2014), similar small growth cone phenotype was observed by expression of constitutively-active, ligand-independent *myr::unc-5* in the VD neurons (Figure 4A-C, and E), consistent with a role of UNC-5 in inhibiting growth cone protrusion.

**Figure 4.**
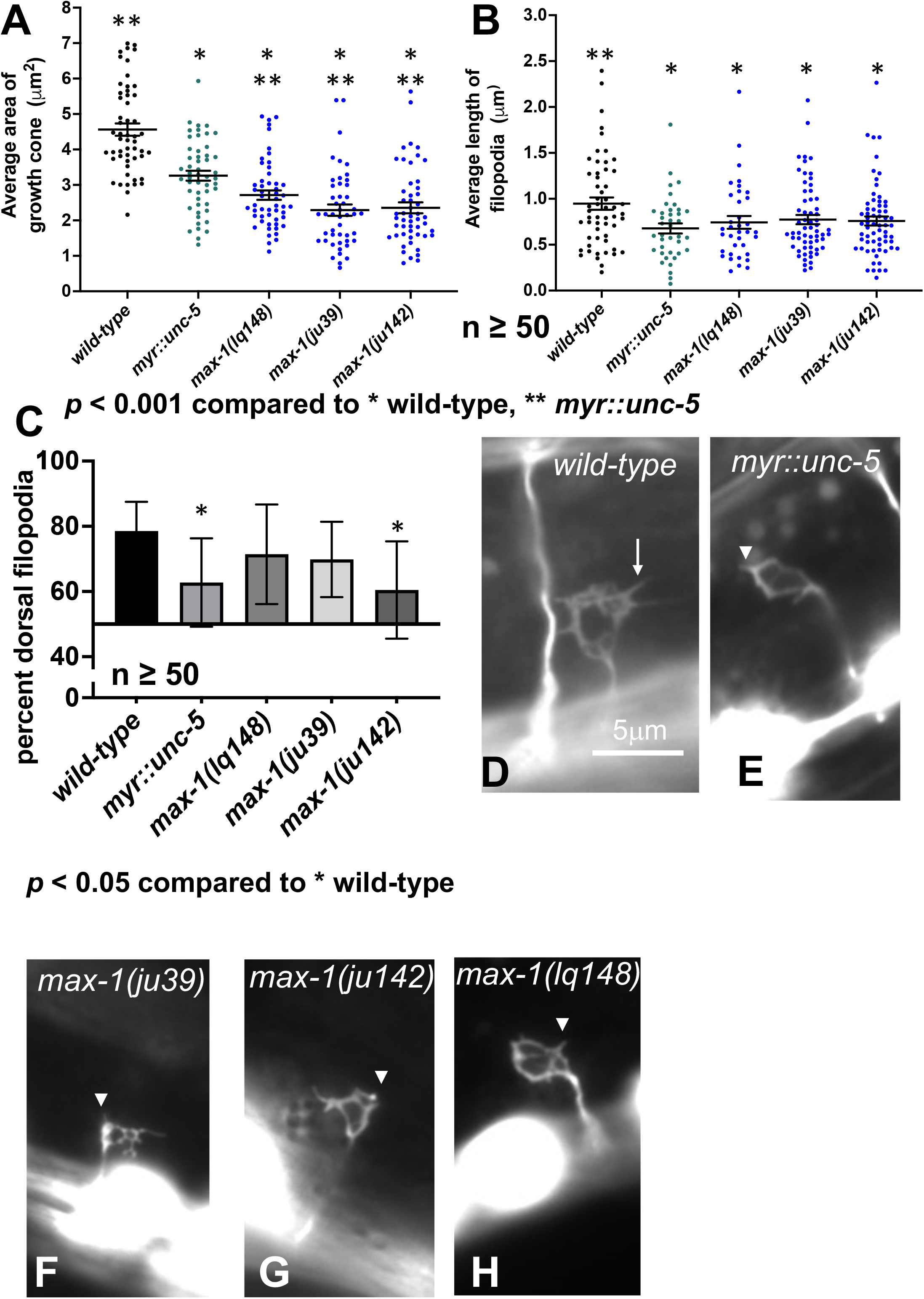
VD growth cone area, filopodia length, and polarity in *max-1* mutants. **(A, B)** Quantification of VD growth cone area (in μm^2^) and filopodial length (in μm) (see materials and methods). Each point represents the score for a single growth cone or filopodium. The mean (line) and standard error of the mean (error bars). significance of difference between genotypes was determined by a two-sided *t-*test with unequal variance. **(C)** A graph showing the percent of dorsally-directed filopodial protrusions in VD growth cones of different genotypes. In wild-type majority of filopodia (78%) extended from the dorsal half of the growth cone. Single asterisks (*) indicates the significant compared to wild-type (*p* < 0.05). Error bar represents 2x standard error of proportion. Significance is determined by Fisher’s exact test. n = number of growth cones. **(D-H)** Fluorescence micrographs of wild-type and mutant VD growth cones. Dorsal is up, anterior is left. The scale bar in D represents 5μm for all micrographs. The arrow in D indicates a filopodial protrusion in wild-type. The arrowheads in E-H point to shortened filopodial protrusions.

### *max-1* suppresses excessive growth cone protrusion in hypomorphic *unc-5* mutants

Our results suggest that the effect of loss of MAX-1 on VD growth cones resembles constitutive UNC-5 activity (small growth cones with reduced protrusion). *unc-5(e152)* and *unc-5(ev480)* are hypomorphic, incomplete loss-of-function mutations with weaker Unc phenotypes than null alleles (Merz *et al*. 2001; Killeen *et al*. 2002). VD/DD axon guidance defects in *unc-5(e152)* and *unc-52(ev480)* were significantly weaker than those in the null allele *unc-5(e791)* (Figure 3). The hypomorphic nature of *unc-5(e152)* and *unc-5(ev480)* suggests that some UNC-5 function is retained in these mutants. VD growth cones in *unc-5(e152)* and *unc-5(ev480)* were more protrusive than wild-type (Figure 5A, B, and D), with increased growth cone area and filopodia length, consistent with the previously-described role of UNC-5 in inhibiting VD growth cone protrusion (Norris and Lundquist 2011; Norris *et al*. 2014; Gujar *et al*. 2018). *unc-5* hypomorph growth cones were also unpolarized (Figure 5C), with loss of dorsal bias of growth cone protrusion as previously reported for *unc-5* mutants.

**Figure 5.**
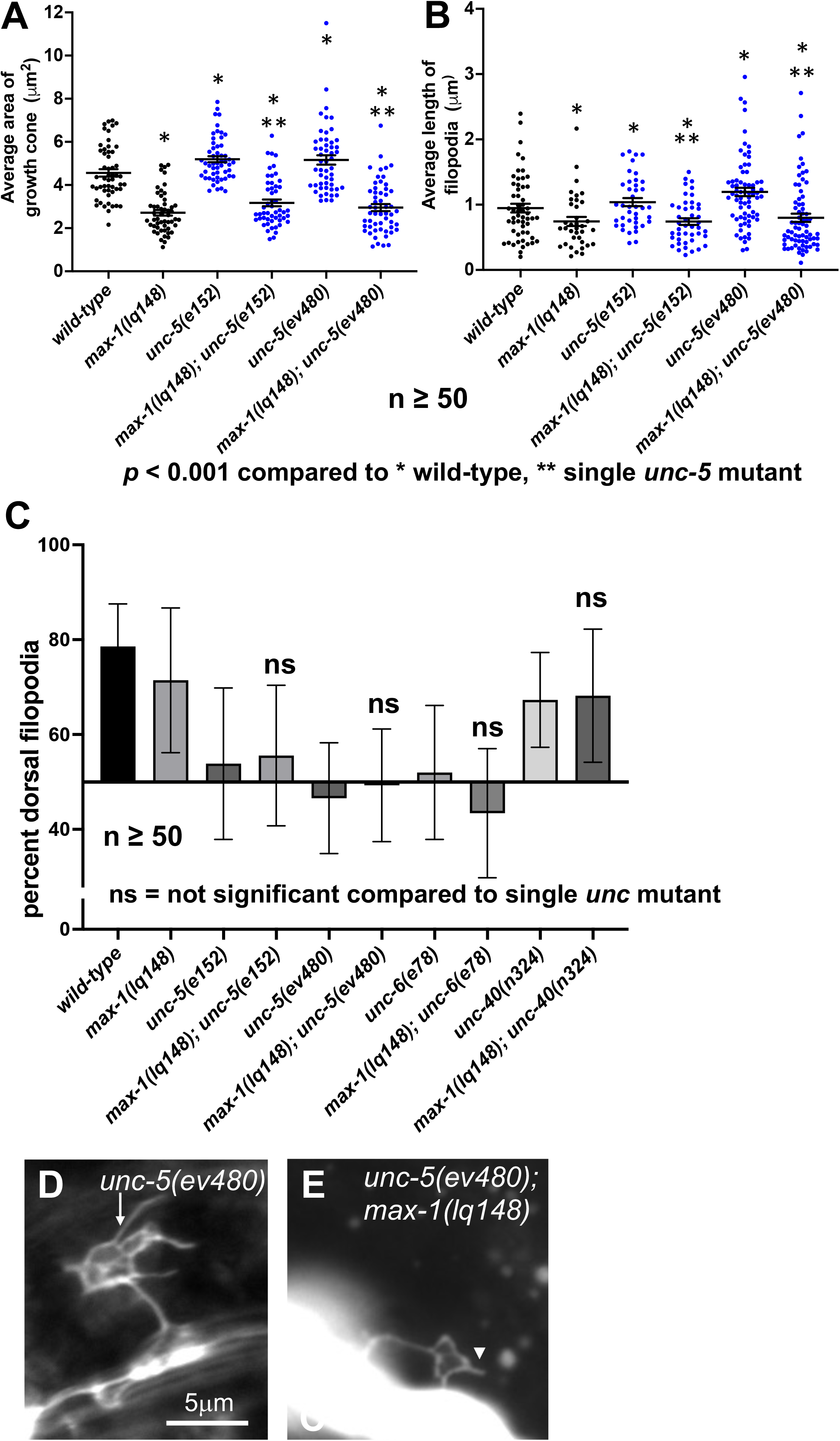
VD growth cone area, filopodia length, and polarity in *max-1; unc-5* double mutants. **(A, B)** Quantification of VD growth cone area and filopodial length as described in Figure 4. **(C)** Quantification of growth cone polarity as described in Figure 4. ns = not significant. **(D, E)** Fluorescence micrographs of mutant VD growth cones as described in Figure 4. The arrow in D indicates an excessively-long growth cone filopodium, and the arrowhead in E indicates a shortened filopodial protrusion.

VD growth cones of *max-1(lq148); unc-5(e152)* and *max-1(lq148); unc-5(ev480)* double mutants displayed reduced protrusion relative to *unc-5(e152)* and *unc-5(ev480)* alone (Figure 5A, B, and E) and in fact resembled the small growth cones of *max-1* alone. Growth cones of *max-1; unc-5(null)* double mutants could not be scored because no VD growth cones emerged from the ventral nerve cord in these animals (data not shown). These results indicate that loss of *max-1* suppressed the excessive VD growth cone protrusion observed in hypomorphic *unc-5* mutants. *max-1* did not suppress *unc-5* growth cone polarization defects, as *max-1; unc-5* growth cones displayed loss of dorsal bias of protrusion as seen in *unc-5* alone (Figure 5C). These data are consistent with the idea that MAX-1 is required for growth cone protrusion but not growth cone polarity, suggested by the *max-1* loss-of-function phenotype.

The UNC-6/Netrin receptor UNC-40/DCC has a dual role in controlling growth cone protrusion. It acts as a heterodimer with UNC-5 to inhibit protrusion, and as a homodimer to stimulate protrusion (Norris and Lundquist 2011; Norris *et al*. 2014; Gujar *et al*. 2018). *unc-40(n324)* null mutants displayed similar growth cone area compared to wild-type, but shortened growth cone filopodial protrusions (Figure 6A-C). This moderate phenotype likely reflects the dual role of UNC-40 in protrusion. *max-1(lq148); unc-40(n324)* double mutant growth cones resembled *max-1(lq148)* alone, with reduced filopodial length similar to *unc-40* and *max-1*, and reduced growth cone area (Figure 6A-C). *unc-40(n324)* growth cones displayed robust dorsal bias of filopodial protrusion similar to wild-type which was not affected by *max-1* (Figure 5C)

**Figure 6.**
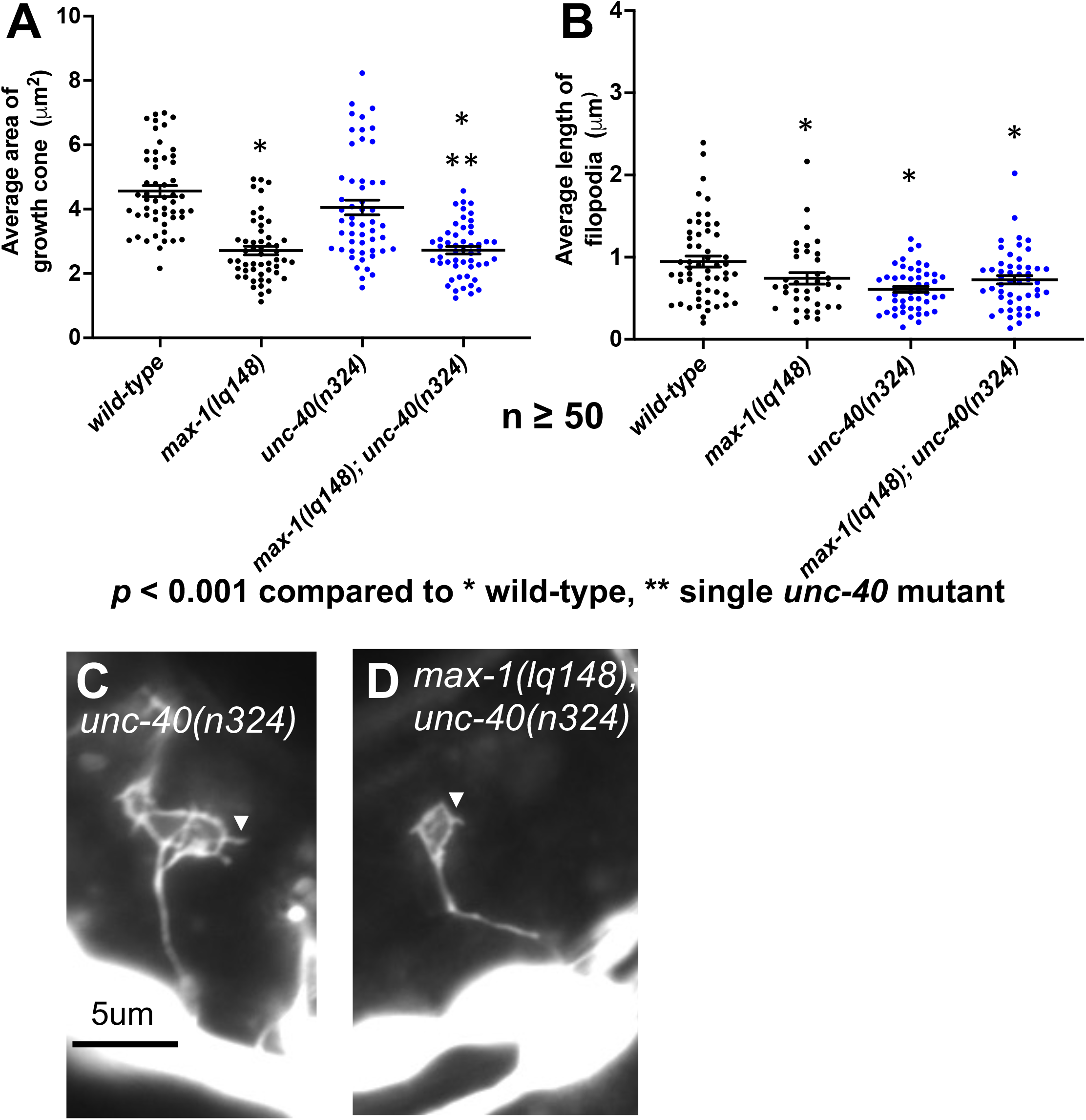
VD growth cone area and filopodia length in *max-1; unc-40* double mutants. **(A, B)** Quantification of VD growth cone area and filopodial length as described in Figure 4. **(C, D)** Fluorescence micrographs of mutant VD growth cones as described in Figure 4. Arrowheads point to filopodial protrusions.

VD growth cones of *max-1; unc-6(ev400)* null double mutants could not be scored because none emerged from the ventral nerve cord. The hypomorphic allele *unc-6(e78)* is a substitution of cysteine to tyrosine in V-3 domain, which disrupts the interaction between UNC-6 and UNC-5 (Lim and Wadsworth 2002; Norris and Lundquist 2011). Dorsal growth is more strongly affected than ventral growth in *unc-6(e78)*. Previous results showed that *unc-6(e78)* mutants displayed unpolarized VD growth cones with excessive protrusion (Norris and Lundquist 2011). When scored for this work, *unc-6(e78)* VD growth cones did not show excessive protrusion (Figure 7A, B, and C), but were still unpolarized as found previously (Figure 5C). *max-1(lq148); unc-6(e78)* VD growth cone protrusion resembled *max-1(lq148)* alone, with reduced growth cone area and filopodia length compared to wild-type and *unc-6(e78)* (Figure 7A, B, and D). *max-1* did not affect loss of growth cone polarity of *unc-6(e78)* (Figure 5C).

**Figure 7.**
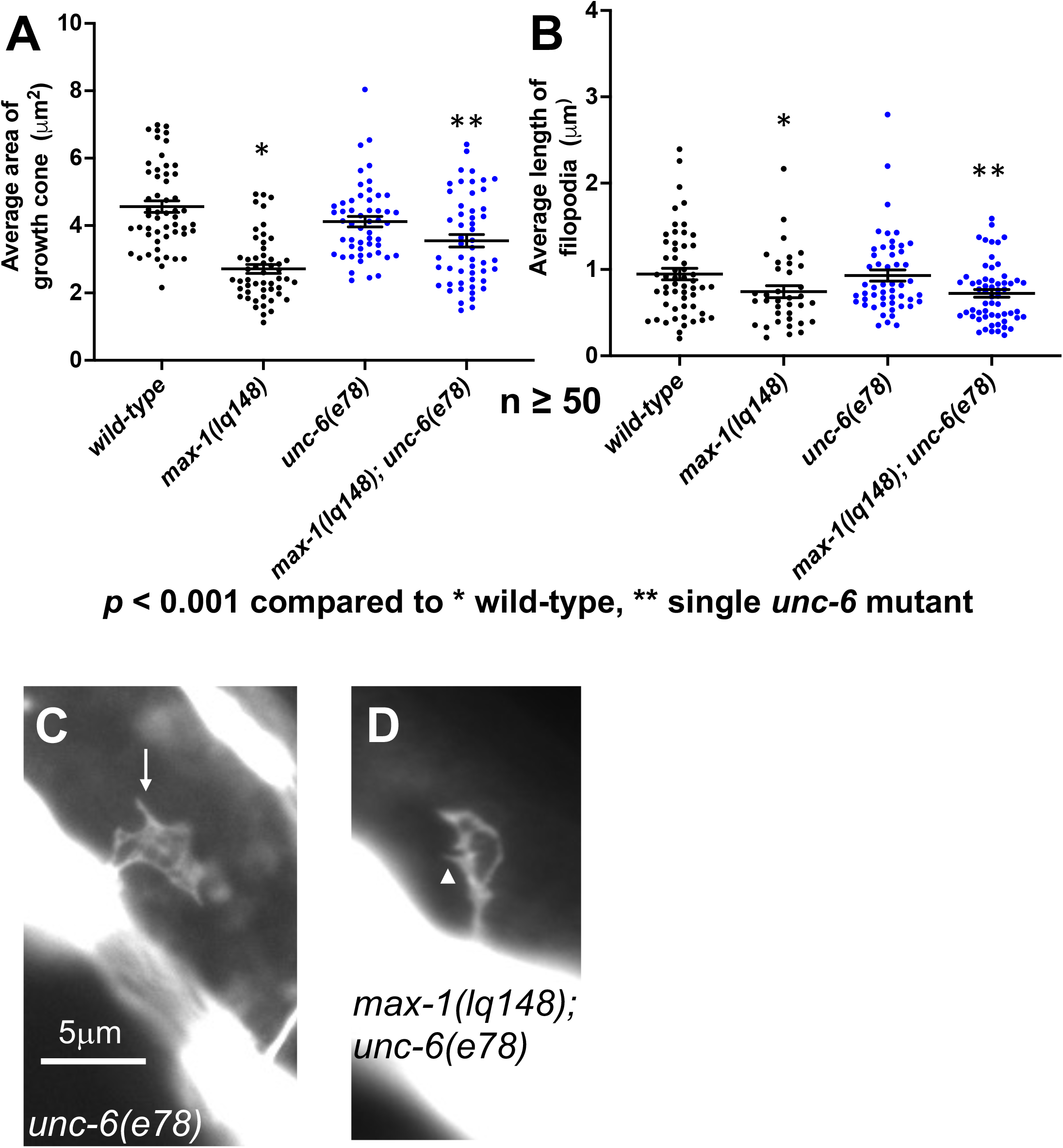
VD growth cone area and filopodia length in *max-1; unc-6* double mutants. **(A, B)** Quantification of VD growth cone area and filopodial length as described in Figure 4. **(C, D)** Fluorescence micrographs of mutant VD growth cones as described in Figure 4. The arrow in C indicates standard-length filopodial protrusion. The arrowhead in D indicates a shortened filopodial protrusion.

### *max-1* enhances VD/DD axon guidance defects of *unc-5, unc-6, unc-40*

Previous studies demonstrated strong genetic interactions between *max-1* and *unc-5, unc-6, and unc-40* in endpoint axon guidance analyses (Huang *et al*. 2002)*. max-1* mutations enhanced hypomorphic *unc-5* alleles, and synergized with *unc-40(null)* for VD/DD axon guidance defects. *unc-5* and *unc-6*, but not *unc-40*, were dominant enhancers of *max-1*. Overexpression of *unc-5* and *unc-6* but not *unc-40* could suppress *max-1*. Finally, overexpression of *max-1* caused axon guidance defects that could be suppressed by overexpression of *unc-5* and *unc-6* but not *unc-40*. Together, these data suggest that MAX-1 acts in the UNC-6 and UNC-5 pathway, possibly in parallel to UNC-40.

We repeated VD/DD axon guidance endpoint analysis using the alleles in which VD growth cones were analyzed here. *max-1* double mutants with *unc-5* and *unc-6* null alleles showed a complete failure of VD/DD axon guidance (Figure 2 and 3), consistent with previous results (HUANG *et al*. 2002) and with our observation of no VD growth cones exiting the ventral nerve cord in these double mutants. *max-1; unc-6(e78)* also showed almost complete axon guidance failure (Figure 2, 3). *max-1* enhanced VD/DD axon guidance defects of the *unc-5(e152)* and *unc-5(ev480)* hypomorphic alleles, with the *max-1; unc-5(ev480)* enhanced synergistically (Figure 2 and 3). This is also consistent with previous results (Huang *et al*. 2002). Axon guidance defects of *unc-40(n324)* were synergistically enhanced by *max-1* (Figure 2 and 3), consistent with previous results (Huang *et al*. 2002). Together, these axon guidance endpoint analyses support previous axon guidance analyses showing that MAX-1 acts in the UNC-5 and UNC-6 pathway, possibly in parallel to UNC-40.

## Discussion

### MAX-1 regulates VD growth cone protrusion, but not polarity

Previous results using VD/DD dorsal axon guidance endpoint analysis indicated that MAX-1 acts with UNC-5 and UNC-6, and in parallel to UNC-40. Our VD/DD axon guidance endpoint analysis presented here supports this conclusion. However, our analysis of VD growth cone morphology during dorsal outgrowth suggests that *max-1* and *unc-5* might have opposing roles. The VD growth cones of *unc-5* hypomorphic mutants displayed excessive protrusion. The growth cones were larger than wild-type with longer filopodial protrusions. This is consistent with the known role of UNC-5 in inhibiting VD growth cone protrusion (Norris and Lundquist 2011; Norris *et al*. 2014; Gujar *et al*. 2018). *max-1* growth cones displayed an opposite phenotype. They were smaller than wild-type with shorter filopodial protrusions, suggesting that MAX-1 was required for robust growth cone protrusion. Thus, in at least some aspects of growth cone outgrowth during axon guidance, UNC-5 and MAX-1 might have opposing roles. MAX-1 might specifically affect growth cone protrusion and not polarity, as *max-1* mutants had only weak polarity defects on their own and did not suppress polarity defects of *unc-5* mutants.

### MAX-1 might attenuate UNC-5 signaling in growth cone protrusion

*unc-5* hypomorphic VD growth cones displayed excessive protrusion, consistent with the role of UNC-5 normally inhibiting growth cone protrusion. However, these hypomorphic alleles retain some UNC-5 activity, as the VD/DD axon guidance defects are significantly less severe than *unc-5* null mutants. VD growth cones of *max-1; unc-5(hypomorphic)* resembled the small, less protrusive growth cones of *max-1* alone. This suggests that MAX-1 is required for the large, overly-protrusive growth cones in *unc-5* mutants. It is possible that MAX-1 acts in a parallel pathway to drive protrusion, such as UNC-40, but genetic analyses suggest that MAX-1 acts in parallel to UNC-40 in the UNC-5 pathway. Rather, we speculate that MAX-1 normally attenuates UNC-5 signaling in growth cone protrusion (Figure 8). In a *max-1* mutant, UNC-5 might be overly-active, leading to reduced growth cone protrusion. In an *unc-5* hypomorph, MAX-1 might be attenuating what little UNC-5 function remains, resulting in large, protrusive growth cones. In a *max-1; unc-5(hypomorphic)* double mutant, *max-1* is no longer attenuating residual UNC-5 activity, which inhibits growth cone protrusion, resulting in suppression of excessive protrusion in *unc-5(hypomorphic)* mutations. If this was the case, we expect that *max-1* would not suppress *unc-5(null)*, because there would be no residual UNC-5 activity. However, we were unable to score VD growth cones in these double mutants, as none emerged from the ventral nerve cord. The phenotypic resemblance of *max-1(null)* mutants and constitutively-active *myr::unc-5* growth cones is consistent with an opposing role of MAX-1 and UNC-5. Furthermore, previous work showing that overexpression of UNC-5 can compensate for overexpression of MAX-1 in VD/DD axon guidance (Huang *et al*. 2002) is consistent with opposing roles, which we discovered in growth cone protrusion.

**Figure 8.**
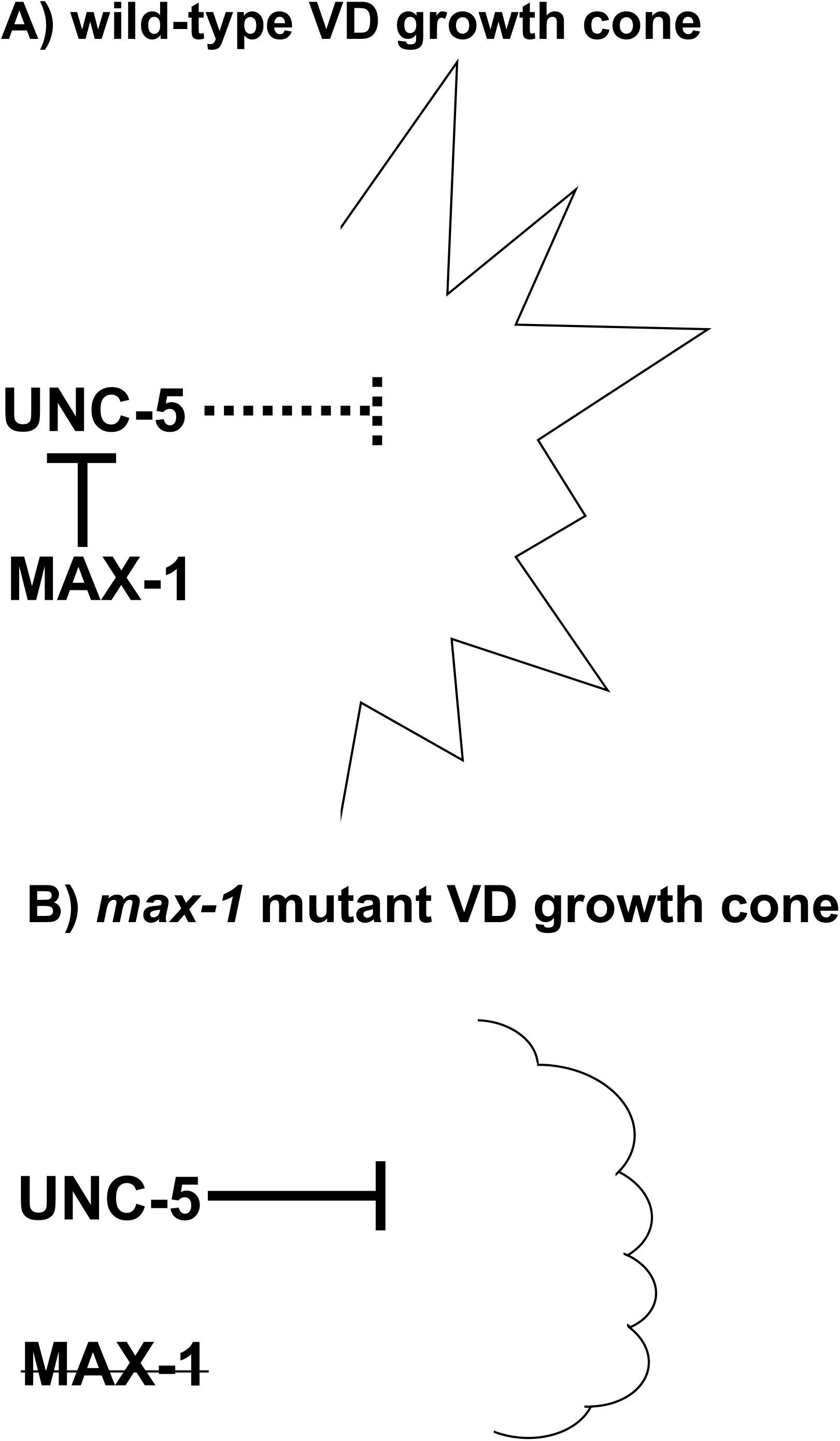
MAX-1 attenuates UNC-5 signaling in VD growth cones. **(A)** In wild-type, MAX-1 normally attenuates UNC-5 signaling, resulting in less inhibition of protrusion and the balanced growth cone protrusion required for proper growth cone outgrowth. **(B)** In *max-1* mutants, UNC-5 is not attenuated and so excessively inhibits protrusion, resulting in small growth cones with limited protrusion. This results in failure of growth cone outgrowth and axon guidance defects.

Previous studies suggest that SUMOylated MAX-1 controls the interaction of the ABP-3/AP-3 complex β molecule with UNC-5, which targets UNC-5 to the lysosome for degradation (Chen *et al*. 2018). In this model, MAX-1 is constitutively bound to UNC-5 until it is SUMOylated, which allows ABP-3 to bind to UNC-5 and target it for lysosomal degradation. This model predicts that loss of MAX-1 would result in reduced UNC-5 activity, yet we show evidence here of increased UNC-5 activity in *max-1* mutants. Possibly, SUMOylated MAX-1 also participates in lysosomal trafficking such that in a *max-1* mutant, UNC-5 is not degraded, resulting in high levels of UNC-5 in the growth cone and inhibited growth cone protrusion.

Too much protrusion, as in an *unc-5* loss-of-function mutant, and too little protrusion, as in *max-1* mutants and constitutively-active *myr::unc-5* animals, both result in qualitatively similar endpoint axon guidance defects. MAX-1 might fine-tune protrusive activity by carefully regulating the activity of UNC-5.

### The relationship of growth cone morphology to endpoint axon guidance phenotypes

Previous results (Huang *et al*. 2002) and results presented here using endpoint axon guidance analysis indicate that MAX-1 and UNC-5 act in the same pathway in a positive manner (i.e. *max-1* and *unc-5* have similar phenotypes that are enhanced in double mutant combinations). Our results analyzing growth cone morphology suggest that MAX-1 and UNC-5 might have opposing roles in growth cone protrusion, with MAX-1 normally attenuating UNC-5 signaling to allow for growth cone protrusion. How these interactions in the growth cone translate to endpoint axon guidance errors remains to be fully elucidated. However, they do point to multiple and complex roles for these molecules in growth cones as they extend. While *max-1* suppresses excessive protrusion in *unc-5(hypomorphic)* mutants, the endpoint axon guidance phenotype is synergistically enhanced in the double mutant. This could be because we only analyzed VD growth cones and effects are different in DD growth cones. More likely is the possibility the roles of these molecules in the growth cone is complex, and that some effects of these mutants on growth cone morphology are not being assayed in this analysis. For example, speed of outgrowth and internal cytoskeletal and endosomal organization was not being assessed here. In any event, these results suggest that in growth cone protrusion, MAX-1 normally inhibits UNC-5 activity, allowing for robust growth cone protrusion necessary for proper VD axon guidance.

### An UNC-6/Netrin-mediated balance of protrusive forces in the growth cone

Previous studies demonstrated that UNC-6/Netrin signaling mediates a balance of protrusive forces in the growth cone. UNC-5 homodimers and UNC-40/UNC-5 heterodimers inhibit growth cone protrusion in response to UNC-6, whereas UNC-40 homodimers stimulate protrusion in response to UNC-6 (Norris and Lundquist 2011; Norris *et al*. 2014; Gujar *et al*. 2018). UNC-5 and UNC-40/UNC-5 inhibit protrusion through the FMO flavin monooxygenases that might destabilize actin and through UNC-33/CRMP, which prevents microtubule + end entry into the growth cone thus preventing entry of pro-protrusive vesicles. The Rac GTPases CED-10 and MIG-2 are involved in both pathways. UNC-40 stimulates protrusion, possibly through the TIAM-1 GEF, the Rac GTPases CED-10 and MIG-2, and actin regulators such as the Arp2/3 complex and UNC-34/Enabled (Norris *et al*. 2009; Norris and Lundquist 2011; Demarco *et al*. 2012). This balance is demonstrated by the observation that the excess growth cone protrusion in *unc-5* mutants requires functional UNC-40 (Norris and Lundquist 2011) (i.e. in an *unc-5* mutant, UNC-40-driven protrusion is not counterbalanced by UNC-5 inhibition of protrusion, resulting in overly-protrusive growth cones).

Work presented here suggests that the PH/MyTH4/FERM myosin-like protein MAX-1 regulates growth cone protrusion by inhibiting UNC-5. *max-1* growth cones are smaller and less protrusive than wild-type, similar to constitutively-activated *myr::unc-5*. Furthermore, *max-1* suppressed the excess growth cone protrusion in *unc-5* hypomorphic mutants, possibly by allowing residual UNC-5 function in these mutants to inhibit protrusion. That too little growth cone protrusion, observed in *max-1* mutants and *myr::unc-5* animals, and too much protrusion, observed in *unc-5* mutants, leads to qualitatively-similar endpoint axon guidance defects underscores the importance of analyzing growth cones during outgrowth to elucidate the molecular and cellular mechanisms that guidance cues and their receptors engage to mediate proper axon guidance and circuit formation.

## Acknowledgements

This work was supported by NIH grants R01040945, R56NS095682, and R03NS114554 to E.A.L. Next generation sequencing was conducted at the KU Genome Sequencing Core, supported by the *Center for Molecular Analysis of Disease Pathways* P20GM103638 (S. Lunte, P.I., E.A.L. co-I.). This work was also supported by the Kansas Infrastructure Network of Biomedical Excellence P20GM103418 (D. Wright, P.I.).

